## Supplemental Figures & Legends for "Synchrony between midbrain gene transcription and dopamine terminal regulation is modulated by chronic alcohol drinking"

**Supplemental Table 1. WGCNA module membership data.** Csv file containing the gene modules identified by WGCNA analysis presented in figure 2.

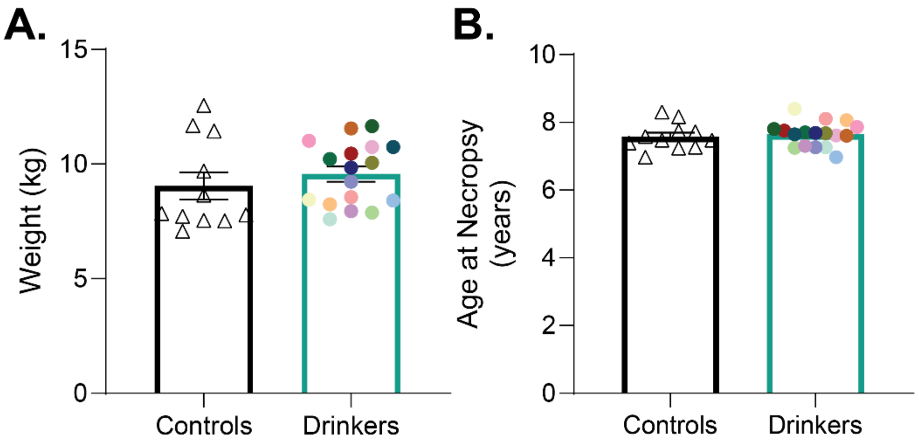


**Supplemental Figure 1. There were no differences in weight or age between the control and drinking subjects at necropsy. (A)** There was no difference in the weight of controls compared to drinkers at the time of necropsy (unpaired t-test; $t_{26} = 0.8255$, $p = 0.4166$). **(B)** There was no difference in age at necropsy between the controls and drinkers (unpaired t-test; $t_{26} = 0.4867$, $p = 0.6306$). (controls: n = 11; drinkers: n = 17) Values indicate mean ± SEM.

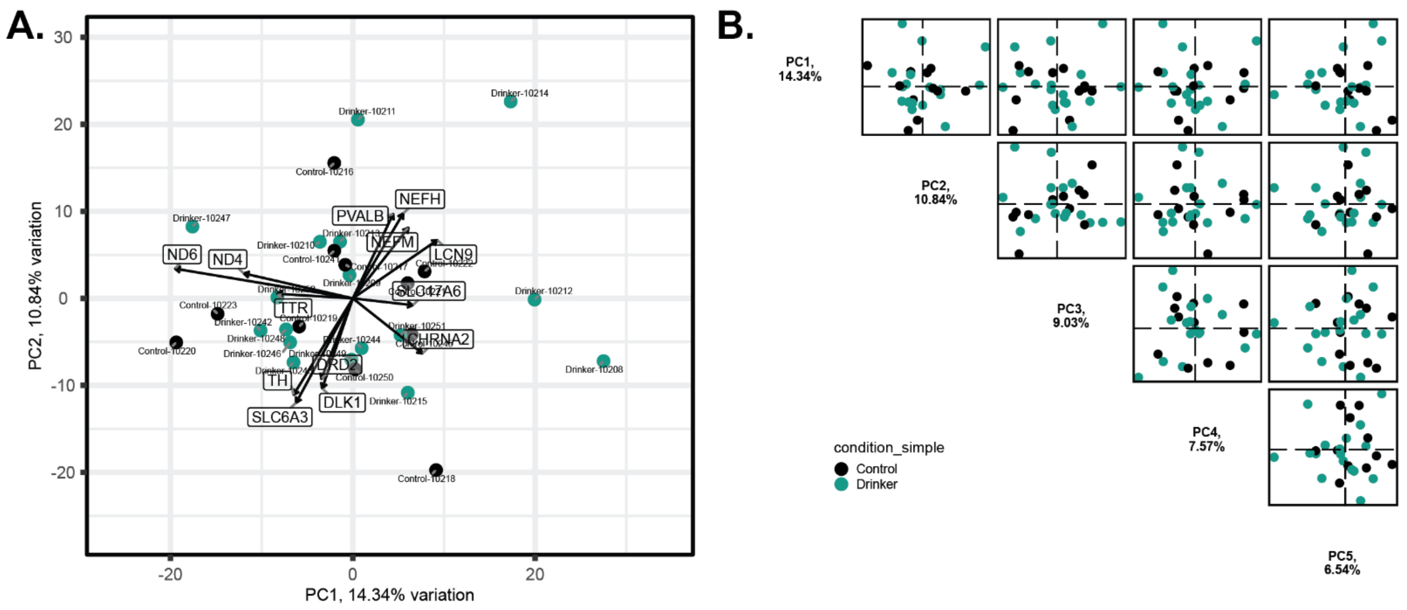


**Supplemental Figure 2. Dimensionality reduction of gene expression via principal component analysis did not produce segregation between drinkers and controls. (A)** There was no appreciable segregation between drinkers and controls across the first 2 principal components. The top genes that influenced the variance are presented along with a vector demonstrating the gene effect through length and the association with other genes through direction. The top genes include genes encoding the dopamine transporter (*SLC6A3*), the D2-type dopamine receptor (*DRD2*), and the rate-limiting enzyme of dopamine synthesis (*TH*). **(B)** Visualization of the top 5 principal components showed no appreciable segregation between groups. (controls: n = 11; drinkers: n = 17)

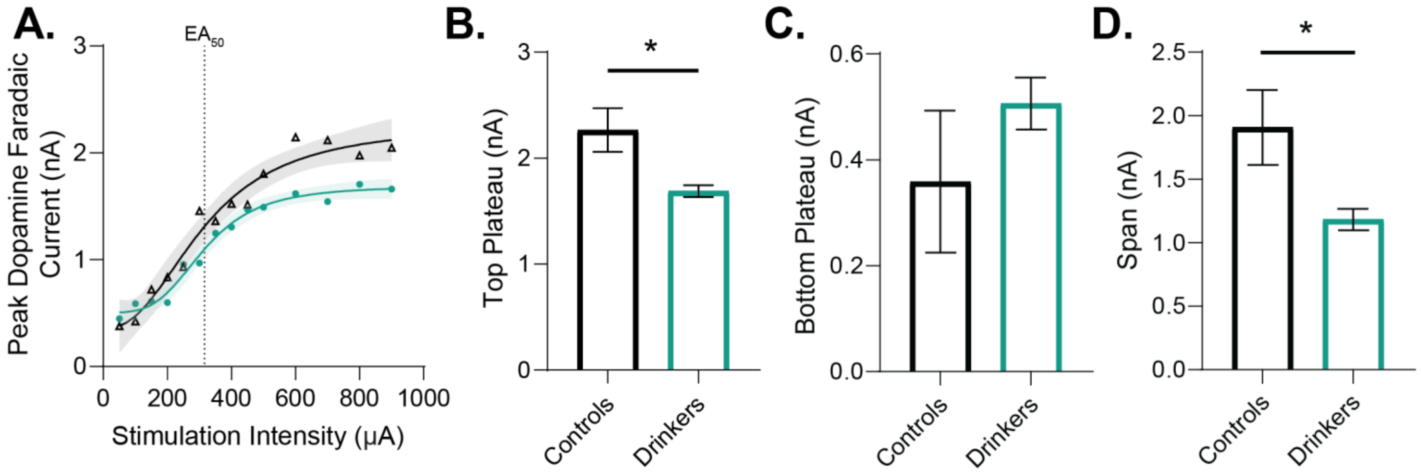


**Supplemental Figure 3. Chronic voluntary alcohol consumption induces long-lasting changes in the dynamics of dopamine release across stimulation intensities. (A)** An input-output curve showing the peak dopamine release (nA) evoked by single pulse stimulations across ascending intensities (50-900 µA). Curves were fit with a 4-parameter sigmoidal regression, and best-fit values are shown with 95% confidence band and half-maximal excitatory amperage ($EA_{50}$) indicated (controls: $EA_{50}$ = 316.4 µA; drinkers: $EA_{50}$ = 315.9 µA). **(B-D)** Comparison of best-fit values between drinkers and controls. **(B)** Subjects with ethanol history have an attenuated upper asymptote compared to controls, indicative of decreased maximal dopamine release magnitude (unpaired t-test; $t_{26}$ = 2.710, p = 0.0118). **(C)** There is no difference between groups at the bottom of the curve plateau (unpaired t-test; $t_{26}$ =1.033, p = 0.3110). **(D)** Drinkers show a decreased span of the input-output curve [upper plateau minus lower plateau] compared to control subjects suggesting a lower dynamic range of dopamine release (unpaired t-test; $t_{26}$ = 2.365, p = 0.0258). (controls: n = 3; drinkers: n = 7) Values indicate mean ± SEM. (* $p \leq 0.05$, ** $p \leq 0.01$, *** $p \leq 0.001$, **** $p \leq 0.0001$)

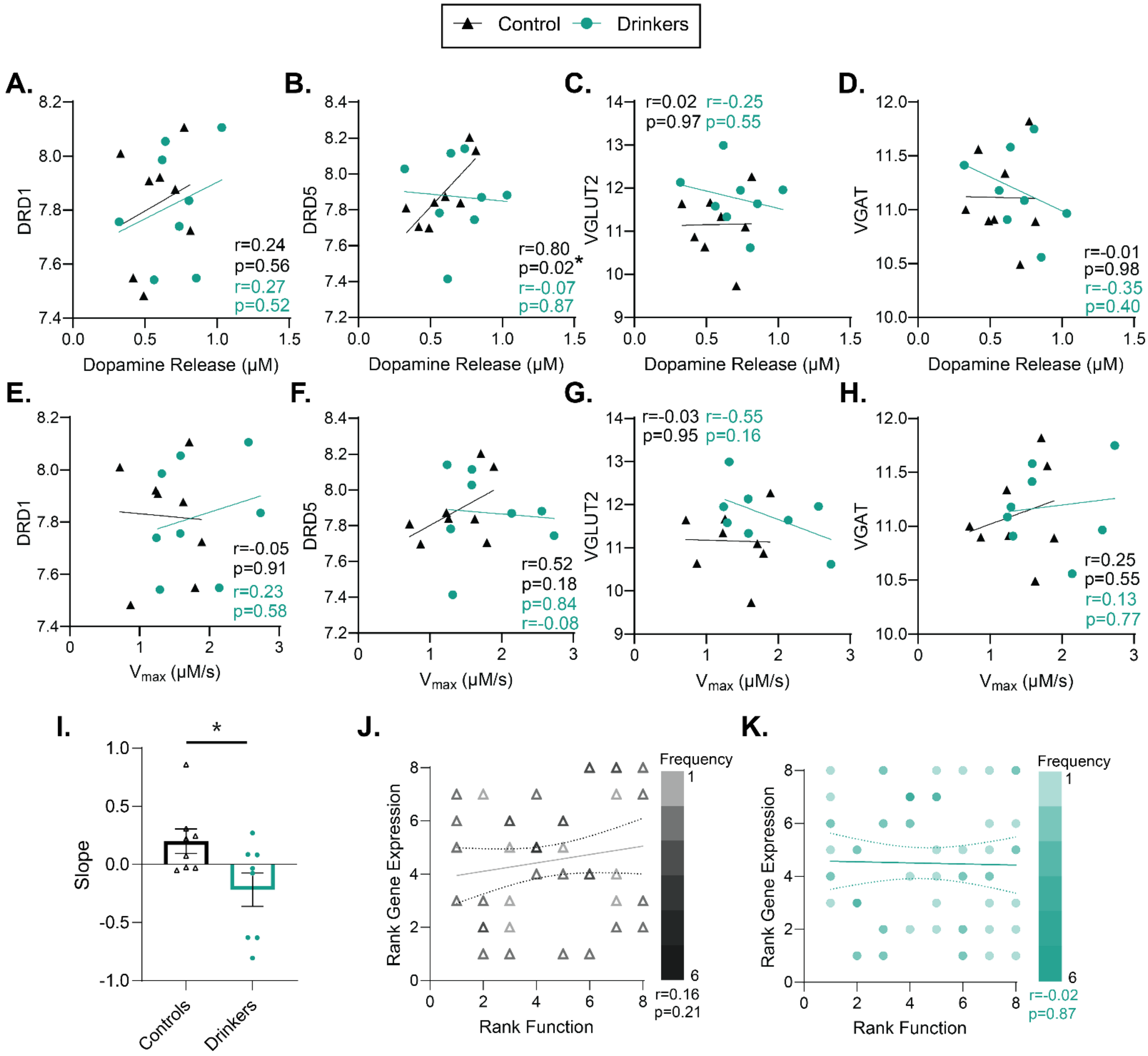


**Supplemental Figure 4. Synchrony between upstream transcription with downstream dopamine terminal function was specific to homosynaptic regulators of the dopamine system. (A-H)** Expression of genes encoding for receptors and transporters in the VTA which are enriched in non-dopaminergic cells was compared to terminal dopamine release and reuptake in the NAc of the same subjects. The best-fit linear regression is plotted for each group and Pearson's correlation coefficient r- and associated p- values are reported as an inset. **(A)** There was no significant correlation between dopamine receptor 1 (*DRD1*) gene expression and dopamine release in either drinkers or controls. **(B)** Dopamine receptor 5 (*DRD5*) gene expression was positively correlated with dopamine release in controls, but not drinkers. **(C)** Expression of vesicular glutamate transporter 2 (*VGLUT2*) did not correlate with dopamine release. **(D)** Vesicular GABA transporter (*VGAT*) did not correlate with dopamine release. **(E-H)** There were no correlations found between $V_{max}$ and expression of *DRD1, DRD5, VGLUT2*, or *VGAT*. **(I)** The mean of the slopes across panels **A-H** were compared between groups and versus zero. While the mean slope was greater in controls than drinkers (unpaired t-test; $t_{14}$ = 2.332, p = 0.0351), neither group differed from 0 (one sample t-test; controls: $t_7$ = 1.884, p

= 0.1016; drinkers: $t_7 = 1.509$, $p = 0.1751$). **(J-K)** To compare the relationship between gene expression and downstream function across transcripts and disparate functional output measures, data were transformed to a within-group rank order for each dependent variable. Thus, each animal was assigned a value from 1 to 8 denoting lowest to highest expression for each of the 4 transcripts, paired with a 1 to 8 value indicating rank order of Vmax and a 1 to 8 value indicating rank order of dopamine release magnitude. This allowed all 8 of correlations above (64 total x-y pairs per group) to be plotted and analyzed in a single coordinate plane with rank-expression on the y-axis and rank-function on the x-axis. Values are presented for each group with the color of the icon indicating the frequency of that coordinate, and the best-fit linear regression is shown with a 95% confidence band. **(J)** There is no correlation between upstream expression of non-dopamine neuron specific receptors and transporters and downstream terminal release dynamics in control subjects. **(K)** There is also no correlation between expression of these genes and accumbal dopamine dynamics after voluntary ethanol consumption and abstinence. Unless otherwise indicated, values indicate mean ± SEM. (controls: n = 8; drinkers: n = 8) (* = $p \le 0.05$, ** = $p \le 0.01$, *** = $p \le 0.001$, **** = $p \le 0.0001$)

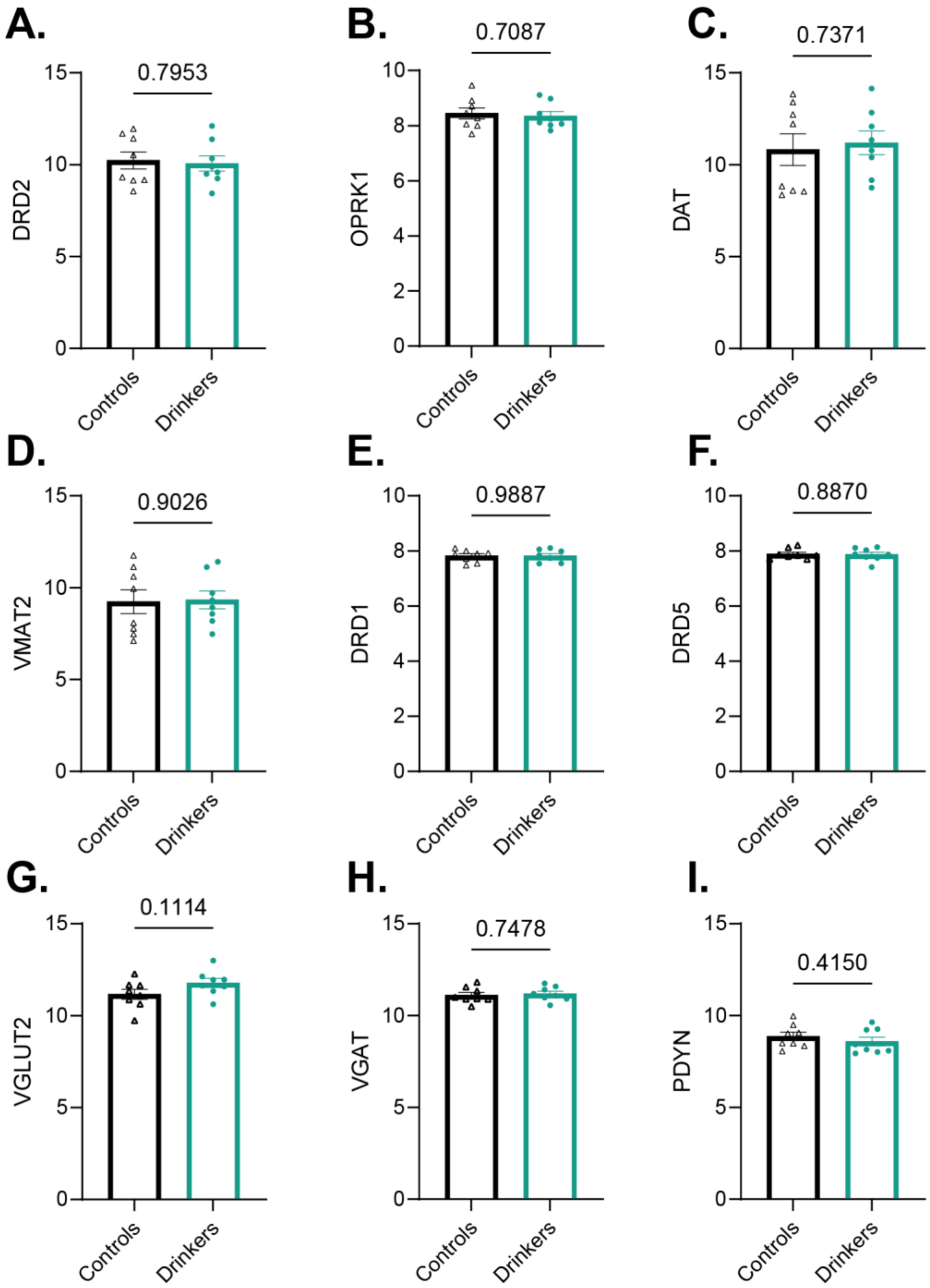


**Supplemental Figure 5. Long-lasting functional changes and changes in transcriptional synchrony occur without significant differences in gene expression between groups.** Drinkers and controls did not differ in VTA gene expression of **(A)** the dopamine receptor 2 (*DRD2*) (unpaired t-test; $t_{14} = 0.2644$, p = 0.7953), **(B)** the kappa opioid receptor (*OPRK1*) (unpaired t-test; $t_{14} = 0.3813$, p = 0.7087), **(C)** the dopamine transporter (*DAT*) (unpaired t-test; $t_{14} = 0.3425$, p = 0.7371), **(D)** the vesicular monoamine transporter 2 (*VMAT2*) (unpaired t-test; $t_{14} = 0.1246$, p = 0.9026), **(E)** the dopamine receptor 1 (*DRD1*) (unpaired t-test; $t_{14} = 0.01468$, p = 0.9887), **(F)** the dopamine receptor 5 (*DRD5*) (unpaired t-test; $t_{14} = 0.1447$, p = 0.8870), **(G)** the vesicular glutamate transporter 2 (*VGLUT2*) (unpaired t-test; $t_{14} = 1.699$, p = 0.1114), **(H)** the vesicular GABA transporter (*VGAT*) (unpaired t-test; $t_{14} =0.3280$, p = 0.7478), **(I)** or prodynorphin (*PDYN*) (unpaired t-test; $t_{14} = 0.8400$, p = 0.4150). (controls: n = 8; drinkers: n = 8) Values indicate mean ± SEM.

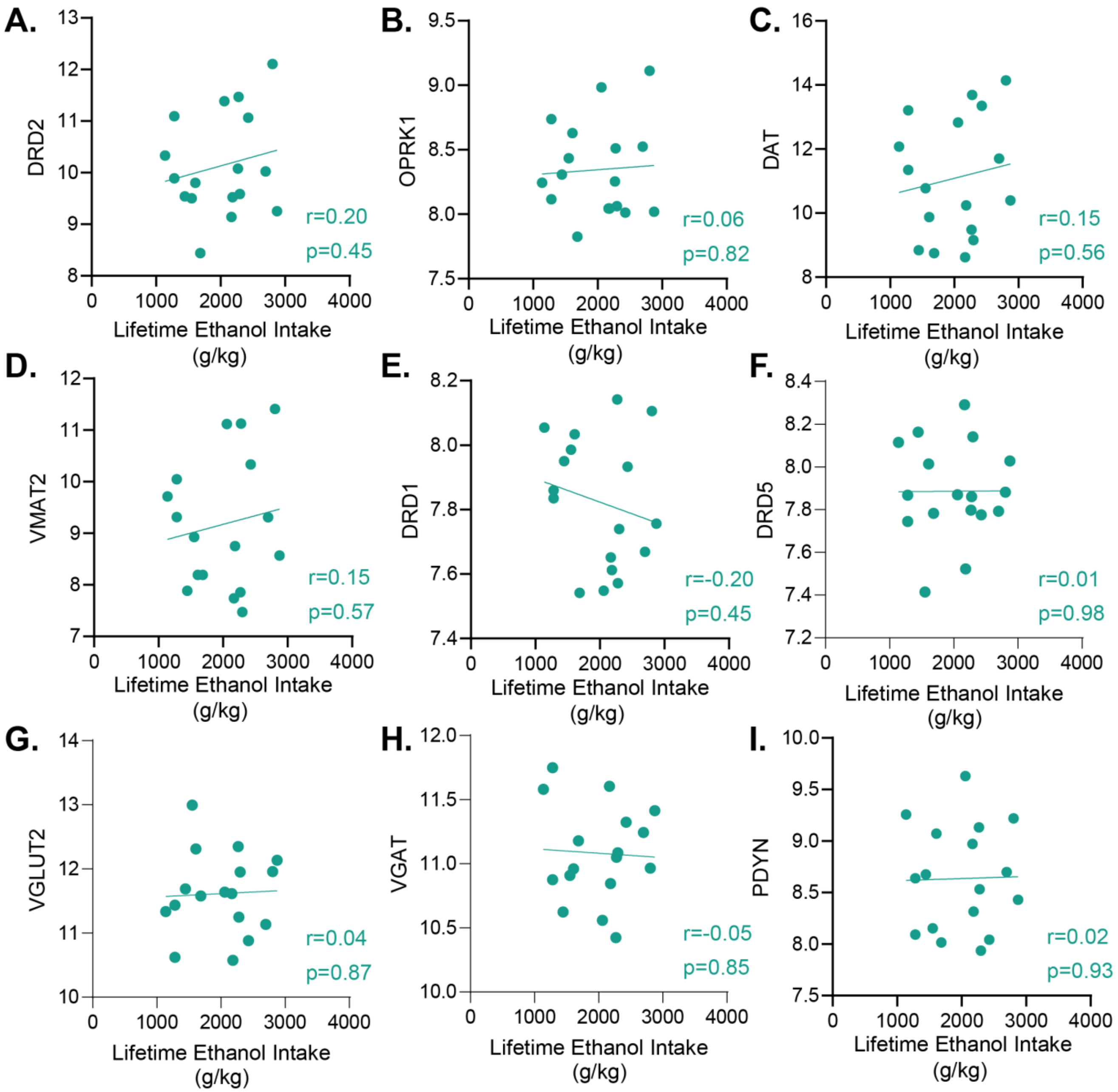


**Supplemental Figure 6. Expression of genes associated with functional changes did not correlate with lifetime ethanol intake.** Lifetime ethanol intake of drinkers did not correlate with gene expression of **(A)** the dopamine receptor 2 (*DRD2*), **(B)** the kappa opioid receptor (*OPRK1*), **(C)** the dopamine transporter (*DAT*), **(D)** the vesicular monoamine transporter 2 (*VMAT2*), **(E)** the dopamine receptor 1 (*DRD1*), **(F)** the dopamine receptor 5 (*DRD5*), **(G)** the vesicular glutamate transporter 2 (*VGLUT2*), **(H)** the vesicular GABA transporter (*VGAT*), **(I)** or prodynorphin (*PDYN*). The best-fit linear regression is plotted for each group and Pearson's correlation coefficient r and associated p values are reported as an inset.

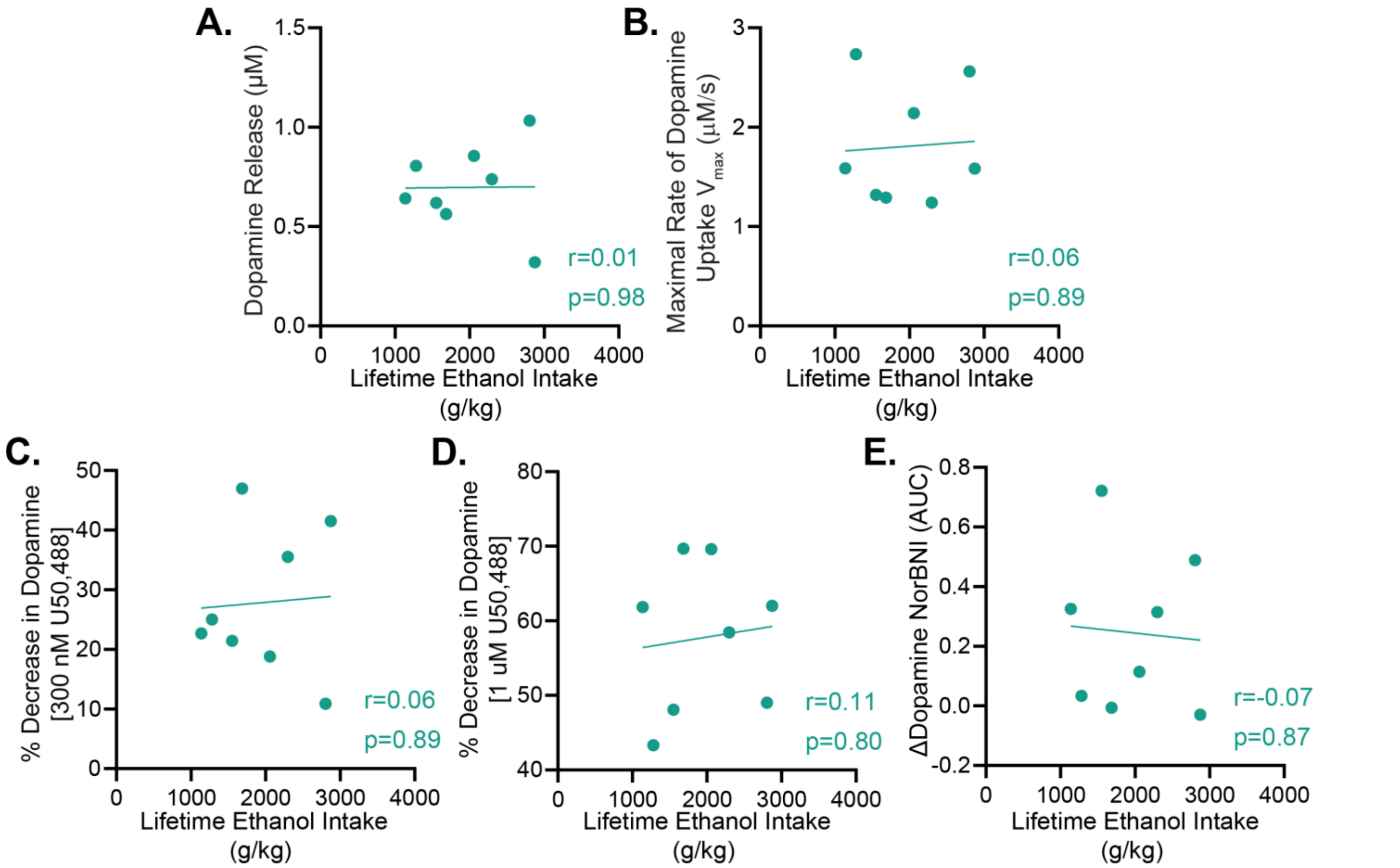


**Supplemental Figure 7. Functional changes at the dopamine terminal did not correlate with lifetime ethanol intake.** Lifetime ethanol intake did not correlate with **(A)** one pulse dopamine release, **(B)** the maximal rate of dopamine reuptake ($V_{max}$) **(C)** the percent decrease in dopamine release with bath application of 300 nM U50,488, **(D)** the percent decrease in dopamine release with bath application of 1 µM U50,488, **(E)** or the percent change in the area under the curve (AUC) of dopamine release with bath application of norbinaltorphimine (NorBNI). The best-fit linear regression is plotted for each group and Pearson's correlation coefficient r- and associated p-values are reported as an inset.

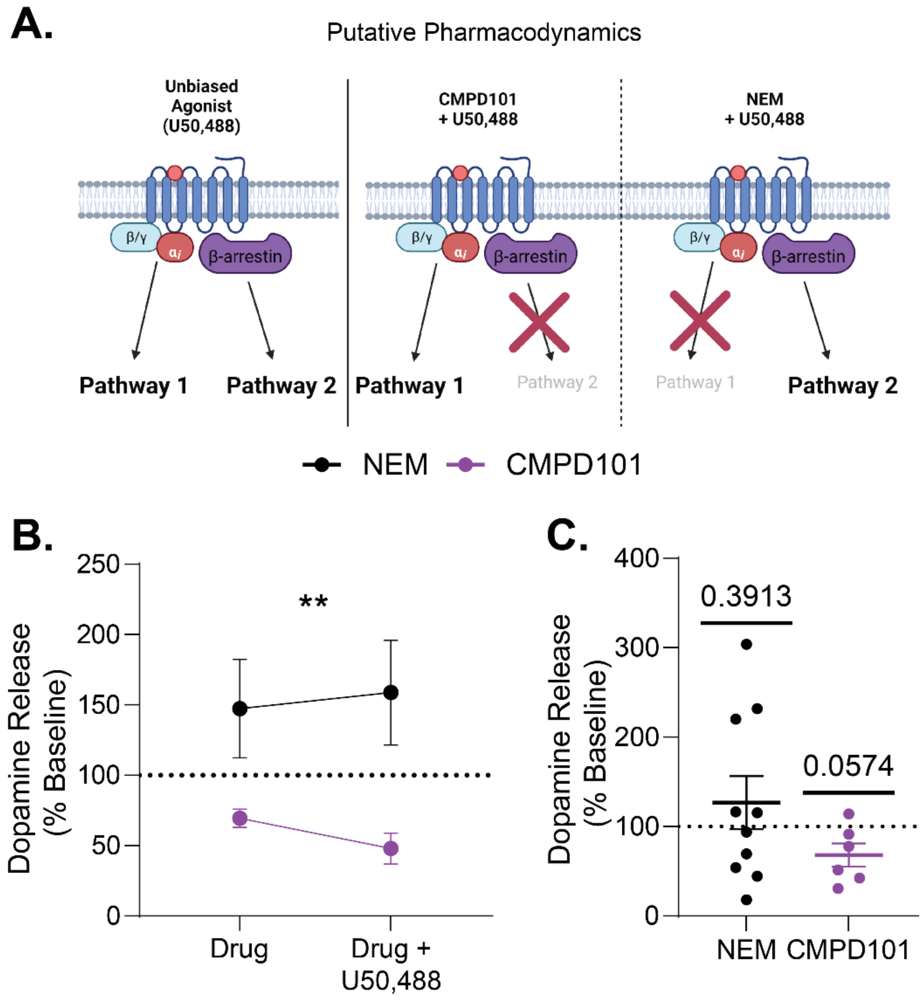


**Supplemental Figure 8. Kappa opioid receptor inhibition of terminal dopamine release was G protein pathway mediated. (A)** Experimental rationale for assessing whether kappa opioid receptor control over dopamine release is mediated by the G protein or β-arrestin pathway. An unbiased agonist, like U50,488, will activate both pathways similarly. Compound 101 (CMPD101) inhibits the β-arrestin pathway, thus, in its presence, application of an agonist would only activate the G protein pathway. N-ethylmaleimide (NEM), in contrast, inhibits G protein signaling, therefore in the presence of an agonist, only the β-arrestin pathway would be activated **(B)** CMPD101 or NEM and U50,488 (a kappa opioid receptor agonist) were applied consecutively and demonstrated differential effects on dopamine release normalized to pre-drug levels (two way ANOVA; wash-on: $F_{1, 26} = 0.02505$, $p = 0.8755$; drug: $F_{1, 26} = 8.645$, $p = 0.0068$; wash-on x drug: $F_{1, 26} = 0.2644$, $p = 0.6114$). **(C)** To account for their different effects, dopamine release after cumulative U50,488 bath application was then normalized to values seen with each of the drugs alone. Application of U50,488 had no effect when NEM was present (one sample t-test; $t_9 = 0.9005$, $p = 0.3913$), but had a trending decrease in dopamine release in the presence of CMPD101 (one sample t-test; $t_5 = 2.458$, $p = 0.0574$) suggesting that G protein activity is necessary to kappa opioid receptor-mediated inhibition of dopamine release. Values indicate mean ± SEM. (* $p \leq 0.05$, ** $p \leq 0.01$, *** $p \leq 0.001$, **** $p \leq 0.0001$)

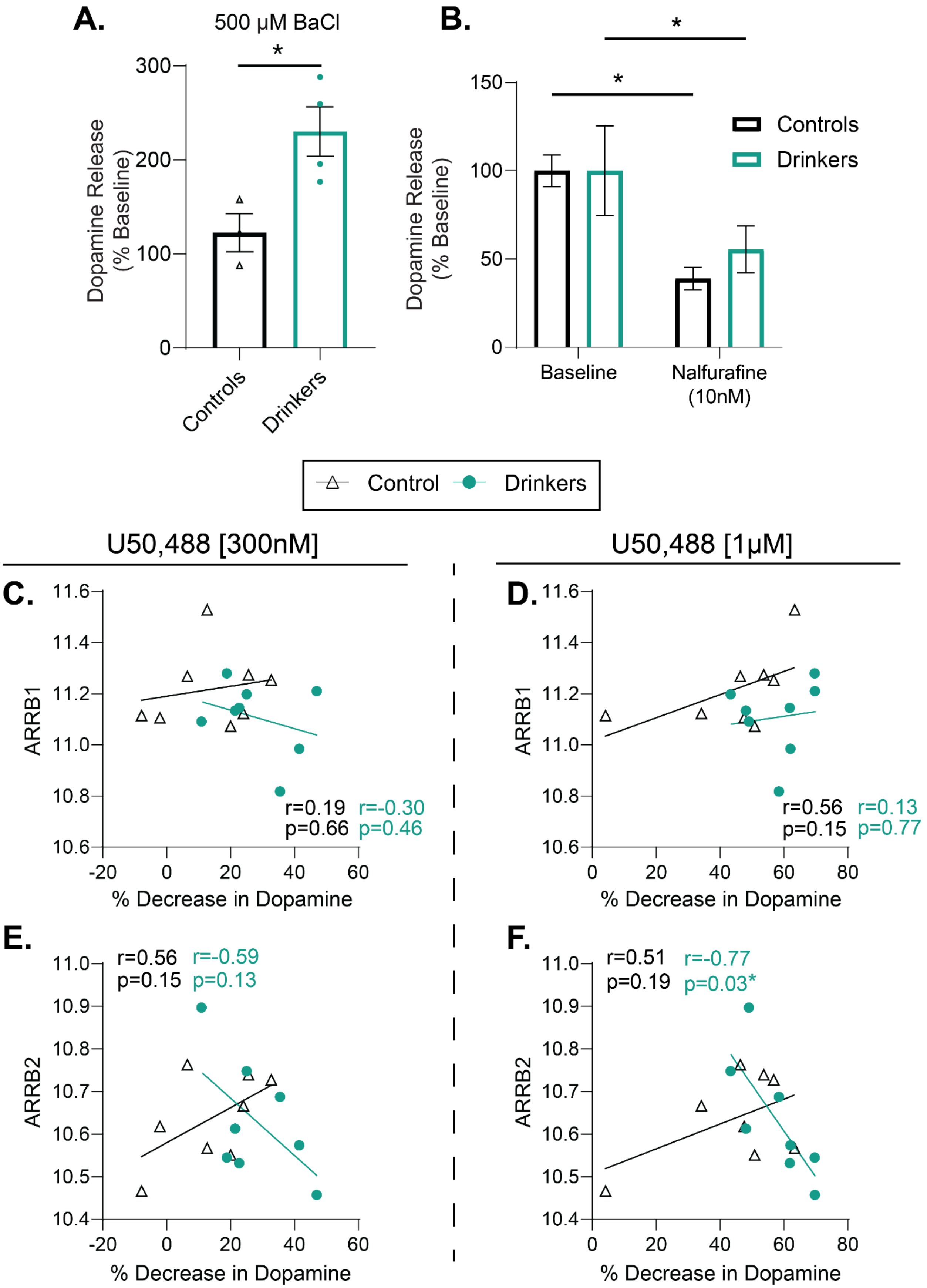
A.
500 µM BaCl
*
Dopamine Release (% Baseline)
300
200
100
0
Controls
Drinkers
B.
*
*
150
100
50
0
Dopamine Release (% Baseline)
Controls
Drinkers
Baseline
Nalfurafine (10nM)
Control
Drinkers
U50,488 [300nM]
U50,488 [1µM]
C.
11.6
11.4
11.2
11.0
10.8
10.6
ARRB1
r=0.19 r=-0.30
p=0.66 p=0.46
-20 0 20 40 60
% Decrease in Dopamine
D.
ARRB1
r=0.56 r=0.13
p=0.15 p=0.77
0 20 40 60 80
% Decrease in Dopamine
E.
r=0.56 r=-0.59
p=0.15 p=0.13
11.0
10.9
10.8
10.7
10.6
10.5
10.4
ARRB2
-20 0 20 40 60
% Decrease in Dopamine
F.
r=0.51 r=-0.77
p=0.19 p=0.03*
ARRB2
0 20 40 60 80
% Decrease in Dopamine

**Supplemental Figure 9. VTA β-arrestin 2 expression was negatively correlated with accumbal kappa opioid receptor inhibition of dopamine release in drinkers. (A)** Application of BaCl, a G protein coupled inwardly rectifying $K^+$ channel inhibitor, increased dopamine release to a greater extent in drinkers compared to controls, suggesting greater levels of endogenous G protein mediated inhibition of dopamine release after ethanol consumption persisting into abstinence (unpaired t-test; $t_5 = 3.407$, $p = 0.0191$). **(B)** Bath application of nalfurafine, a G protein biased kappa opioid receptor agonist, decreased dopamine release in both drinkers and controls (two way ANOVA; concentration: $F_{1, 6} = 30.76$, $p = 0.0015$; group: $F_{1, 6} = 0.1010$, $p = 0.7614$; concentration x group: $F_{1, 6} = 0.7562$, $p = 0.4180$; Šídák's multiple comparisons test: baseline vs. nalfurafine; controls: $t_6 = 4.058$, $p = 0.0133$; drinkers: $t_6 = 3.818$, $p = 0.0175$). **(C-F)** VTA expression of *ARRB1* (β-arrestin 1 / arrestin-2) and *ARRB2* (β-arrestin 2 / arrestin-3) were correlated with the percent decrease of baseline dopamine release observed at 300 nM and 1 μM of the kappa opioid receptor agonist U50,488. **(C)** *ARRB1* expression was not associated with the effect of 300 nM U50,488 in decreasing dopamine release in either group. **(D)** The effect of kappa opioid receptor activation by 1 μM U50,488 was not associated with *ARRB1* expression in either group. **(E)** At the low concentration, *ARRB2* expression was not correlated with inhibition of dopamine release in either group, though the slope of the best fit linear regression is negative in drinkers. **(F)** At the higher dose there was no association between the efficacy of U50,488 and expression of *ARRB2* in controls. However, in drinkers there was a negative correlation such that greater efficacy of U50,488 at the kappa opioid receptor was associated with decreased *ARRB2* expression. Best fit linear regression is shown with Pearson’s correlation coefficient r and associated p values are reported as an inset. Values indicate mean ± SEM. (* $p \le 0.05$, ** $p \le 0.01$, *** $p \le 0.001$, **** $p \le 0.0001$)

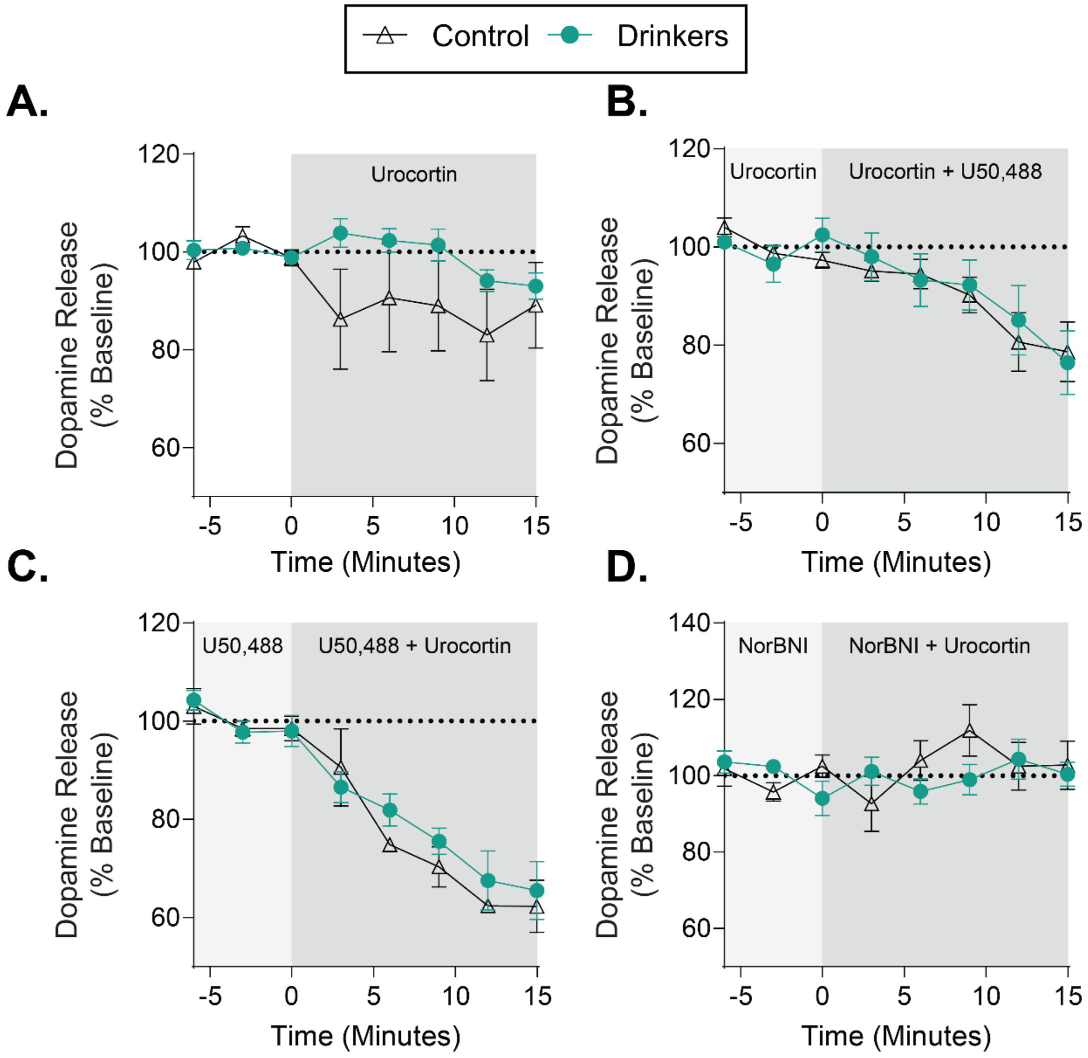


**Supplemental Figure 10. Corticotrophin releasing factor (CRF) receptor-mediated inhibition of dopamine release requires kappa opioid receptor activation. (A)** Urocortin, a CRF receptor agonist, had no effect on accumbal dopamine release in drinkers or controls. (two way ANOVA; time: $F_{1.572, 22.01} = 2.340$, $p = 0.1290$; group: $F_{1, 14} = 1.358$, $p = 0.2633$; time x group: $F_{7, 98} = 1.525$, $p = 0.1678$). **(B)** Consecutive bath application of U50,488, a kappa opioid receptor agonist, decreased dopamine release in both groups similarly, demonstrating a loss of supersensitization of the receptor in the presence of CRF receptor activation (two way ANOVA; time: $F_{2.266, 31.72} = 11.67$, $p < 0.0001$; group: $F_{1, 14} = 0.03315$, $p = 0.8581$; time x group: $F_{7, 98} = 0.4261$, $p = 0.8838$). **(C)** When a kappa opioid receptor agonist was present, the addition of urocortin further decreased dopamine release in drinkers and controls (mixed effects analysis; time: $F_{2.923, 28.39} = 35.53$, $p < 0.0001$; group: $F_{1, 10} = 0.4990$, $p = 0.4961$; time x group: $F_{7, 68} = 0.5462$, $p = 0.7964$)**. (D)** Urocortin had no effect on dopamine release when applied in the presence of a kappa opioid receptor antagonist, norbinaltorphimine (NorBNI) (mixed effects analysis; time: $F_{7, 72} = 0.8132$, $p = 0.5793$; group: $F_{1, 11} = 0.2384$, $p = 0.6349$; time x group: $F_{7, 72} = 1.438$, $p = 0.2037$). Values indicate mean ± SEM. (controls: n = 8; drinkers: n = 8)
